# Spontaneous retrotranspositions in normal tissues are rare and associated with cell-type-specific differentiation

**DOI:** 10.1101/536896

**Authors:** Xiao Dong, Lei Zhang, Kristina Brazhnik, Moonsook Lee, Xiaoxiao Hao, Alexander Y. Maslov, Zhengdong Zhang, Tao Wang, Jan Vijg

## Abstract

Activation of retrotransposons and their insertions into new genomic locations, i.e., retrotranspositions (RTs), have been identified in about 50% of tumors. However, the landscape of RTs in different, normal somatic cell types in humans remains largely unknown. Using single-cell whole-genome sequencing we identified 528 RT events, including LINE-1 (L1), and Alu, in 164 single cells and clones of fibroblasts, neurons, B lymphocytes, hepatocytes and liver stem cells, of 29 healthy human subjects aged from 0 to 106 years. The frequency of RTs was found to vary from <1 on average per cell in primary fibroblasts to 7.8 per cell in hepatocytes. Somewhat surprisingly, RT frequency does not increase with age, which is in contrast to other types of spontaneous mutation. RTs were found significantly more likely to insert in or close to target genes of the Polycomb Repressive Complex 2 (PRC2), which represses most of the genes encoding developmental regulators through H3K27me3 histone modification in embryonic stem cells. Indeed, when directly comparing RT frequency between differentiated liver hepatocytes with liver stem cells, the latter were almost devoid of RTs. These results indicate that spontaneous RTs are associated with cellular differentiation and occur, possibly, as a consequence of the transient chromatin transition of differentiation-specific genes from a transcriptionally repressed to activated state during the differentiation process.

---

Retrotransposons are widespread repetitive elements in the genome. They are usually classified into LTR (Long Terminal Repeat) retrotransposons and non-LTR retrotransposons. The latter type is the most abundant and include Long Interspersed Nuclear Element (LINE)-1 (L1), Alu and SVA elements. Together these account for more than one-third of the human genome^1^. Only L1 elements are autonomous retrotransposons and about 100 of them have been demonstrated as still active in the human genome^2-4^. Most are inactivated by truncations, rearrangements and other mutations. L1 elements can be activated, transcribed and, after reverse transcription, re-integrated in the genome^2,5^. While normally repressed, possibly through epigenetic mechanisms^6^, retrotranspositions (RTs) have been reported in both tumors and in the germline^4,7-9^.

In normal somatic cells, derepression of retrotransposons has been documented during early embryonic development, neurological disorders, and replicative senescence^10-13^. Activation of retrotransposons has been implicated in the loss of genome integrity during aging of somatic cells^14^. However, the landscape of RTs in normal human cells is almost completely unknown apart from some conflicting reports on human neurons^15-18^. Indeed, quantitative information about RTs cannot be obtained by studying bulk tissues and requires advanced, well-validated single-cell genomics technology^19,20^.

To identify RTs in normal somatic cells, we used whole-genome sequencing data of 152 single cells and 12 single-cell derived clones, of 56 B lymphocytes, 28 fibroblasts, 36 neurons, 36 hepatocytes and 8 liver stem cells, of 29 healthy subjects aged between 0 year and 106 years. Apart from our own data this also included single-cell whole genome sequences generated by others using different single-cell amplification methods (Table S1)^20-22^. Whole-genome sequencing of bulk DNA of the same subjects were also analyzed to filter out germline polymorphisms. On average, 30.3±8.4x depth of single cells and clones, and 29.4±7.2x depth of bulk DNA were obtained after alignment, covering on average 87.3±9.2% and 91.9±0.3% of the genome of single cells/clones, and bulk DNA separately (Table S2). Somatic RTs were then identified comparing the alignment of single cells and clones to their corresponding bulk DNAs using TraFiC^8,9^. We validated our variant calling by PCR analysis of 10 randomly chosen RTs (Supplementary Information; Table S3). In addition, to ensure that none of the identified RTs were artifacts of the amplification we directly compared RT frequencies obtained after whole-genome amplification of single human fibroblasts with those found in unamplified DNA from clones derived from single cells in the same population of cells (Fig. S1). The results obtained show very similar results between single-cell and unamplified clone analysis, indicating that the RTs identified were bona fide insertions and not artifacts of the amplification procedure or variant calling.

Across all samples we identified 528 RT events. Consistent with previous studies in tumors^7-9^, most of the RTs were L1 insertions (77.5%), with Alu as the second most frequent type (14.2%) (Fig. 1a, Fig. S2a,b, Table S4). RT frequency appeared to be highly cell-type specific, with almost no insertions found in fibroblasts, 2-3 per cell in B lymphocytes and neurons and more than 7 in hepatocytes (Fig. 1b, Fig. S3); about 50% to 90% of the cells carried at least one RT depending on the cell-type (Fig. 1c, Fig. S4a,b). The RT frequency observed is in agreement with a previous report of almost no RTs in fibroblast clones ^23^. The frequency observed in neurons is largely in agreement with ref ^16^, which reported <1 RT per neuron, but not in agreement with ref ^15^, which reported 13.7 RTs per neurons.

**Fig. 1.**
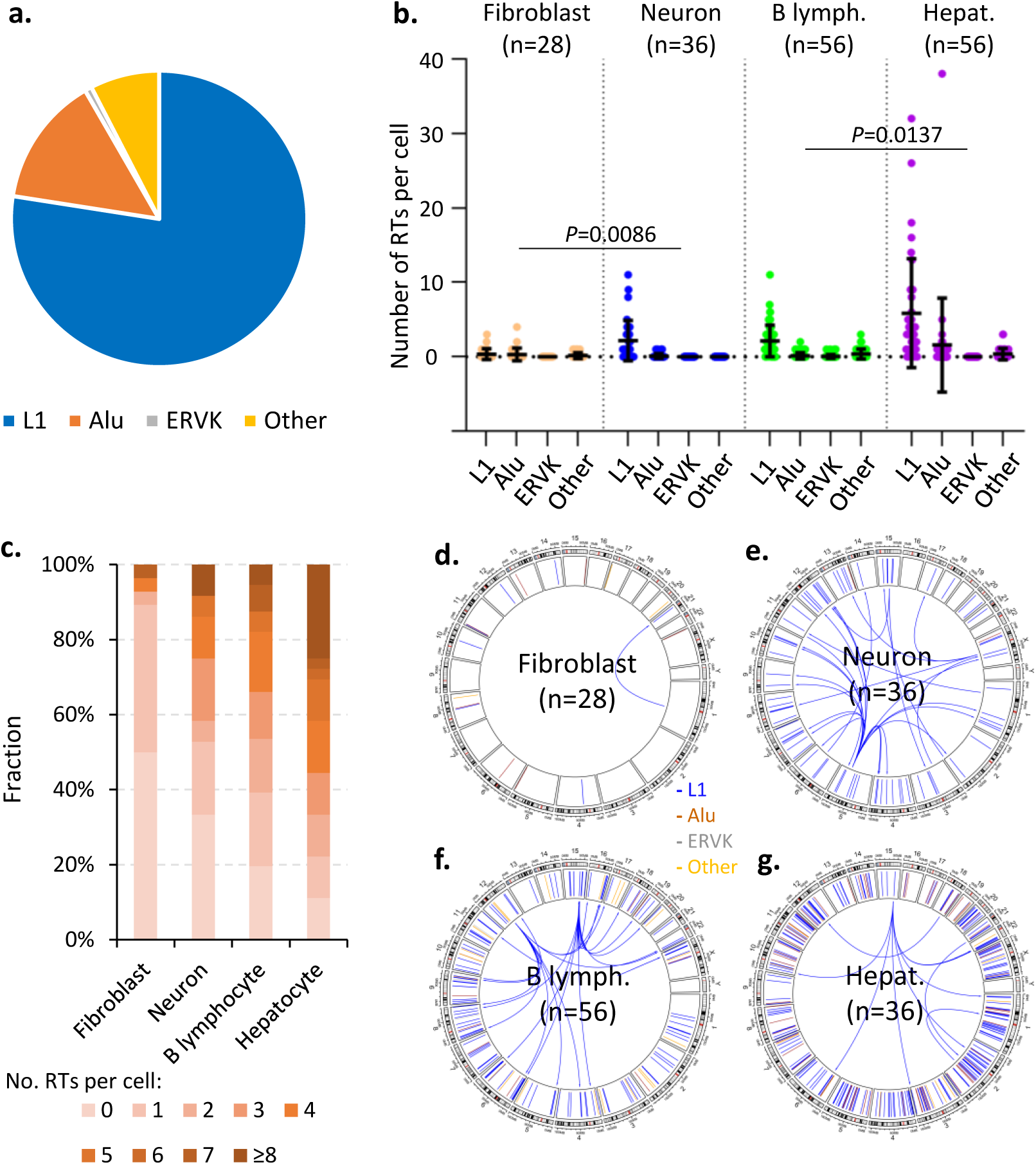
RT spectrum, frequency and distribution. (**a**) Spectrum of RTs identified across all cells collectively. (**b**) Frequencies of RTs in each cell type. A dot presents a cell; bars present the average with its standard deviation (s.d.). *P* values were determined for the difference in total RTs between cell types using two tailed Student’s t-test. (**c**) Fractions of cells with different numbers of RTs. (**d**-**g**) Genome distribution of RTs in each cell type. RTs of all cells per cell type were plotted together in each figure. Within each circus plot, links in the inner layer present directions from L1 sources to their insertion sites. Only sources of those with transductions can be determined by aligning non-repetitive sequences transduced with RT to the reference genome (Supplementary Information), and were plotted. The middle layer of a circus plot presents insertion sites of all RTs. The external layer presents cytobands of chromosomes.

For the most frequent L1 family, germline RT occurs from a small number of polymorphic, “hot” elements^4^. We identified three hot L1 source elements, which were also found to be cell-type specific (Fig. 1d-g, Table S5): 15q24.3 is the source of 19 events in 8 hepatocytes and 11 B lymphocytes of 14 human subjects; 5q23.1 is the source of 17 events in 5 neurons of 2 subjects; and 12q14.3 is the source of 10 events in 3 B lymphocytes of 2 subjects. Although similar numbers of hot L1 sources were reported in cancers^9^, there is virtually no overlap with the L1 sources in normal cells. Indeed, Xp22.2-Xp22.13, a hot L1 source found in cancers, accounted for only one single event in a B lymphocyte. This cell-type specificity in source elements may be a result of potential differences in transcriptional activation of L1 sources between tumors and normal cells.

Thus far, only expression but not re-integration of retrotransposons has been studied as a function of age. The wide age-coverage of the samples of three cell types, i.e., B lymphocytes, hepatocytes and neurons, allowed us for the first time to test if RT insertions accumulate during aging. To our surprise, unlike other types of mutations ^24-26^, no increase in RT frequency was observed in any of the three cell types, B lymphocytes (aged between 0 and 106 years), hepatocytes (5 months −77 years), and neurons (15 −42 years) during aging (Fig. 2a, Fig. S5). This is in spite of the demonstrated increase in expression of retrotransposons during aging^27-29^. This finding suggests that RT activation may be a transient event occurring only once during a cell’s lifetime after which repression is again implemented. To test this, we studied possible preferences in genomic location of the RTs identified.

**Fig. 2.**
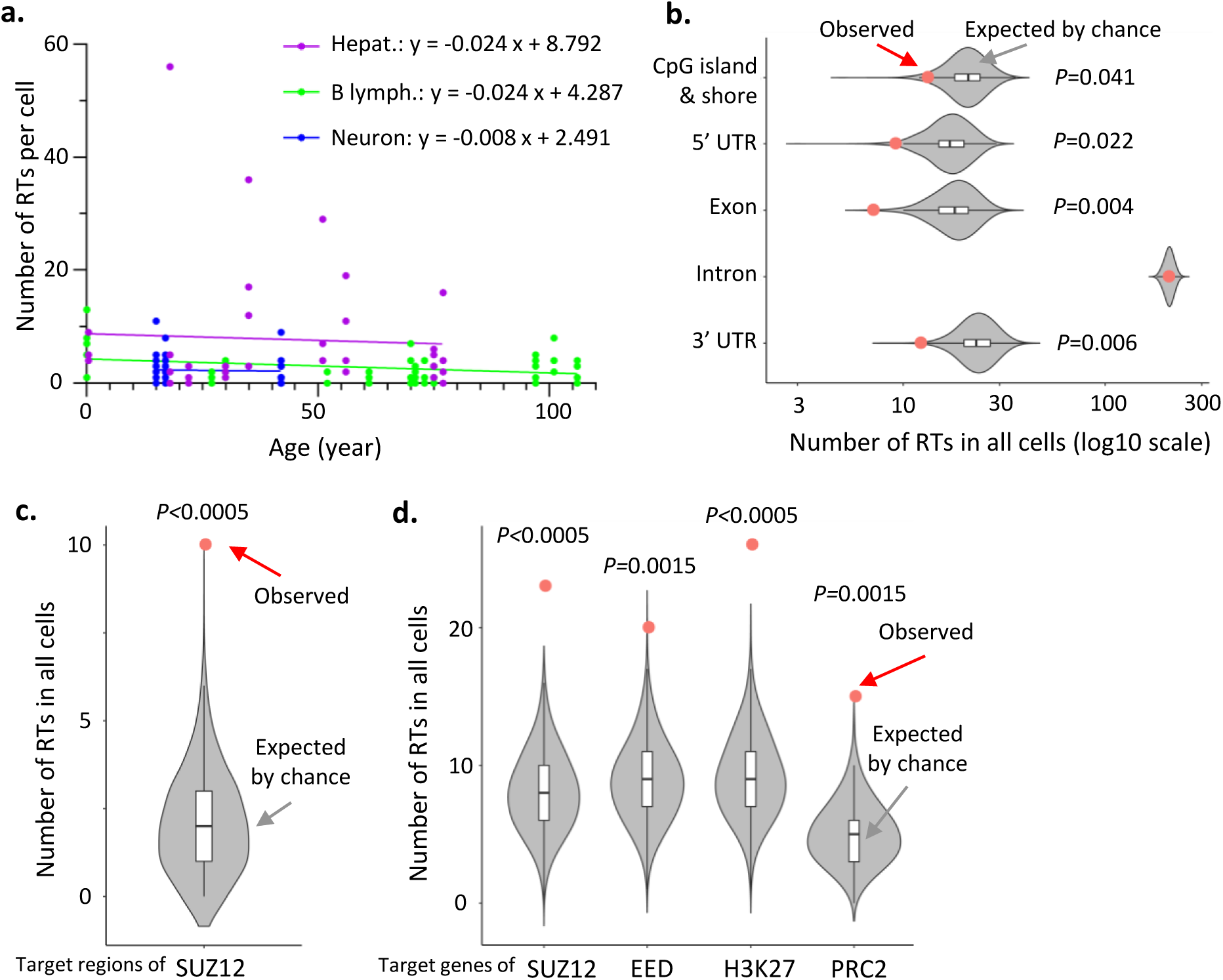
Age-related frequency and genomic-feature distribution of RTs. (**a**) Frequency of RTs during aging. Linear regressions were performed for each cell type separately. (**b**) Depletion of RTs in genomic features. (**c**) Enrichment of RTs in target regions of SUZ12. (**d**) Enrichment of RTs in target genes of PRC2. (**b**-**d**) A red dot presents the number of RTs observed. A violin plot (with a box plot inside) presents a distribution (with median and quantiles) of expected numbers of RTs by chance alone (2,000 times of random sampling, Supplementary Information). *P* values were estimated as Monte Carlo *P* values based on the random sampling.

RT distribution across the genome was analyzed for all RTs collectively. The results showed that insertion sites are significantly depleted from the most obvious functional sequences. That is, the RT frequencies in gene exons, 5’ and 3’ UTRs are 62%, 48%, and 49%, respectively, significantly less than what would be expected by chance alone (*P*=0.004, 0.022 and 0.006, permutation test; Fig. 2b, Fig. S6a-c). RT insertion sites are also depleted from CpG islands and their flanking island shores (38% less than the frequency expected by chance alone; *P*=0.041), which is consistent with the finding that insertions have preferentially been found at AT regions in tumors^7^.

We then tested the target regions of 161 transcription factors (TFs), identified from multiple cell types by ENCODE^30^, for enrichment or depletion of RTs. For most of the TFs, their target regions were neither depleted nor enriched for RTs (Table S6). However, the target regions of seven and three TFs were found depleted or enriched for RT insertions, respectively (fold change≥2 and FDR-adjusted *P*<0.05 for multiple testing correction; Fig. 2c, Fig. S7). Interestingly, the most significant association was observed as an enrichment of RTs at the target regions of SUZ12, i.e., 4.8-fold higher frequency than expected by chance alone (*P*<0.0005 permutation test, FDR-adjusted *P*=0.0345).SUZ12 is a major component of the Polycomb Repressive Complex (PRC2), which functions in embryonic stem (ES) cells to repress expression of developmental genes^31^. The majority of genes encoding developmental regulators exhibit extended regions of PRC2 binding in humans ^32^.

We then validated the enrichment at PRC2 target regions with an independent dataset of ES cells^32^, in which target genes of PRC2 were defined as the intersection of three target gene sets of SUZ12, EED and H3K27me3 separately. We found that RT insertion sites are significantly enriched in any of the three gene sets as well as their intersection, i.e., the PRC2 target genes, (3.0-fold higher than expected by chance alone, *P*=0.0015 permutation test; Fig. 2d, Fig. S8a-c, Table S7).

Based on the observed enrichment of RT insertions at PRC2 and H3K2me3 target genes, we hypothesized that these events are in some way associated with the temporal regulation of chromatin during differentiation from a stem or progenitor cell to a fully mature cell type. To test this, we made use of the available whole-genome sequences of both differentiated, mature liver hepatocytes and liver stem cells. The identity of the latter was verified using a set of stem-cell-specific and epithelial-progenitor-cell-specific cell surface markers (EpCAM, Lgr5, CD90, CD29, CD105, and CD73; Supplementary Information). If RT would be a transient event occurring only once in a lifetime, we would expect no RTs in the stem cells. Indeed, we found that although hepatocytes have the highest RT frequency (7.83±11.51 RTs per cell, avg.±s.d.) among the four cell types analyzed, liver stem cells have almost no RTs, i.e., 0.25±0.48 per cell with 25% of cells carrying at least one insertion (*P*=0.00037, Fig. 3a,b, Fig. S9). This result confirmed that RTs are associated, at least in human liver, with cellular differentiation.

**Fig. 3.**
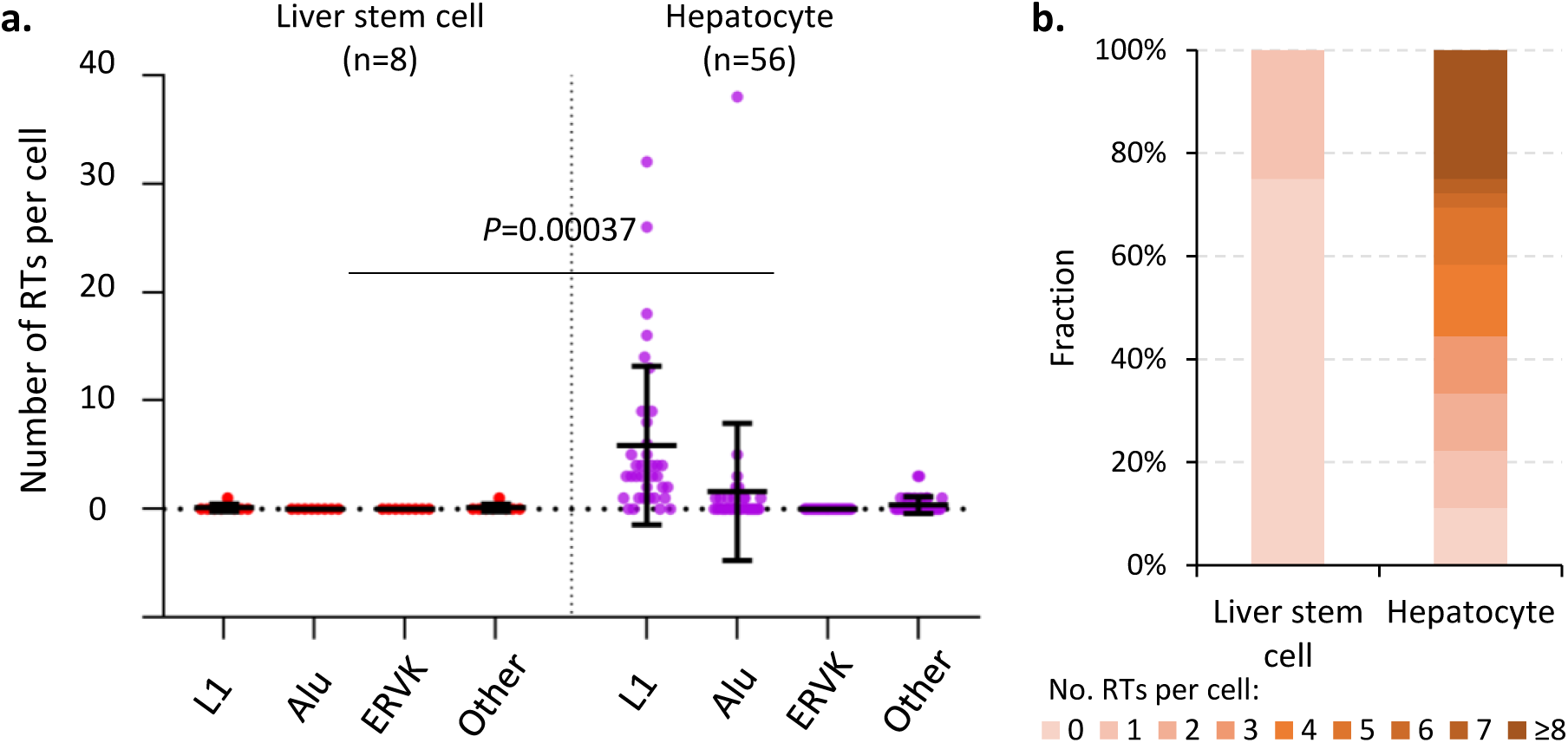
RT frequency in liver stem cells and hepatocytes. (**a**) Frequencies of RTs in liver stem cells and terminally differentiated hepatocytes. A dot presents a cell; bars present the average and s.d.. *P* values were determined for the difference of total RTs using two tailed Student’s t-test. (**b**) Fractions of cells with different number of RTs.

The cell type-specific differences observed could well be related to the state of differentiation and/or the origin of the cell type. For example, fibroblasts are the least specialized cells in the connective-tissue family ^33^, which may be related to their very low frequency of RTs. The RTs during differentiation may have both beneficial and damaging effects. As documented, expression of certain L1 elements is probably required during early embryonic development^34,35^. However, L1 insertions are stochastic events and of low abundance, i.e., no more than a few per cell. Therefore, it seems highly unlikely that they have any consistent regulatory effect on development. Instead, they are probably mere by-products of essential biological processes.

In summary, using single-cell whole-genome sequencing, we demonstrate that RTs in normal somatic cells of healthy human subjects occur at a low frequency, which is cell type-specific. Multiple lines of evidence indicate that rather than gradually increasing with age, for example, as a consequence of an age-related activation ^27^, spontaneous RTs in normal cells occur mainly during cellular differentiation, probably as by-products of the dramatic chromatin alterations associated with terminal differentiation. While this suggests that RTs do not significantly contribute to the normal aging process due to their low frequency and lack of further accumulation, RTs may well play a role in age-related diseases, most notably cancer ^7-9,18^.

## Acknowledgements

This study was supported by NIH grants P01 AG017242 (J.V.), K99 AG056656 (X.D.), P01 AG047200 (J.V.), U01 ES029519 (J.V.) and the Paul F. Glenn Center for the Biology of Human Aging.

## Author Contributions

J.V. and X.D. conceived this study and designed the experiments. X.D., X.H., T.W. and Z.Z. analyzed the data. L.Z., K.B., M.L. and A.Y.M. performed the experiments. X.D. and J.V. wrote the manuscript.

---

**Fig. S1. Comparison of RT frequency observed in single cells and single-cell derived clones.**

Clones and SCMDA-amplified single cells were obtained from the same population of human fibroblasts^20^; in the paper about LIANTI amplification, single cells of another human fibroblast population were used and amplified using multiple different amplification protocols^22^. No significant difference was observed in number of RTs per cell between the three datasets.

**Fig. S2. Spectra of RTs in different cell types.**

(**a**) Spectrum of RTs in all cells collectively. L1-td0, td1, and td2 represent solo-L1 events, partnered transductions and orphan transductions, respectively (Supplementary Information). (**b**) Spectra of RTs in different cell types.

**Fig. S3. Average frequencies of RTs in each cell type.**

L1-td0, td1, td2 represent solo-L1 events, partnered transductions and orphan transductions, respectively (Supplementary Information).

**Fig S4. Fractions of cells with different numbers of RTs.**

(**a**) L1. (**b**) Alu.

**Fig S5. Frequency of L1 insertions during aging.**

Linear regressions were performed for each cell type separately.

**Fig. S6. Depletion of RTs in genomic features per cell type.**

A red dot represents the number of RTs observed. A violin plot (with a box plot inside) presents the distribution (with median and quantiles) of expected numbers of RTs by chance alone (2,000 times of random sampling, Supplementary Information). *P* values were estimated as Monte Carlo *P* values based on the random sampling.

**Fig S7. Enrichment of RTs in target regions of TFs.**

A red dot represents the number of RTs observed. A violin plot (with a box plot inside) presents the distribution (with median and quantiles) of expected numbers of RTs by chance alone (2,000 times of random sampling, Supplementary Information). *P* values were estimated as Monte Carlo *P* values based on the random sampling.

**Fig S8. Enrichment of RTs in target genes of PRC2.**

A red dot represents the number of RTs observed. A violin plot (with a box plot inside) presents the distribution (with median and quantiles) of expected numbers of RTs by chance alone (2,000 times of random sampling, Supplementary Information). *P* values were estimated as Monte Carlo *P* values based on the random sampling.

**Fig S9. Average frequencies of RTs in liver stem cells and hepatocytes.**

L1-td0, td1, and td2 represent solo-L1 events, partnered transductions and orphan transductions, respectively (Supplementary Information).

