## Supplementary Text for "Spontaneous retrotranspositions in normal tissues are rare and associated with cell-type-specific differentiation"

### Sample collection

The experimental procedures in this study were approved by the Einstein-Montefiore institutional review board (IRB). Besides samples collected at the Albert Einstein College of Medicine, the others were obtained as described below. Cord blood of two human subjects were obtained from AllCells Inc. and StemCell Technologies Inc. separately. Hepatocytes of eight human subjects were obtained from Lonza Walkersville Inc. Liver stem cells of one human subject were obtained from Kerafast, Inc.

### Single-cell isolation

For B lymphocytes, PBMCs were isolated from whole blood and bulk B lymphocytes were isolated using MACS separation (Kit #130-050301, Miltenyi Biotec) by the Molecular Cytogenetics Core at the Albert Einstein College of Medicine. Single B lymphocytes were isolated using the CellRaft system (Cell Microsystems) following the same protocol as described previously[1], except an additional step, i.e., coating the array with gelatin (2% in water; G1393-100ML Sigma-Aldrich) at 37°C for one hour to help the B lymphocytes attaching after initial rinsing of the array[2,3]. Single B lymphocytes were collected into PCR tubes with 2.5 µl PBS and frozen immediately on dry ice and kept at -80°C until use.

For hepatocytes, single hepatocytes with diploid genomes were isolated using fluorescent activated cell sorting (FACS) with DNA-binding dye Hoechst 33342 (Life Technologies) and LIVE/DEAD Cell Vitality Assay Kit $C_{12}$ Resazurin/SYTOX™ Green (Thermo Fisher Scientific). Single hepatocytes were collected into PCR tubes with 2.5 µl PBS and frozen immediately on dry ice and kept at -80°C until use.

Single fibroblasts were isolated as described previously[1].

### Single-cell whole-genome amplification

Single-cell whole-genome amplification of the isolated single cells was performed using the single-cell multiple-displacement amplification (SCMDA) procedure that we developed previously[1]. The amplicons were selected using an 8-locus locus-dropout test as described[1]. The selected amplicons were used for library preparation and sequencing.

**Preparing single-cell derived clones**

Liver stem cells from a one-year-old human subject were obtained from Kerafast, Inc. The cells were cultured in polarization media (DMEM, 10% dialyzed FBS (Invitrogen), 1.5mM Xanthosine (Sigma), 1x Penicillin/Streptomycin, 20ng/ml EGF human (Invitrogen), 0.5ng/ml TGF beta human recombinant (Sigma)) according to the manufacturer's protocol (Kerafast, Inc.) [4-6]. To verify liver stem cell identity we used a set of stem-cell-specific and epithelial-progenitor-cell-specific cell surface markers, i.e., EpCAM, Lgr5, CD90, CD29, CD105, CD73, using flow cytometry analysis (FACS; LSRII, Becton Dickenson) as recommended previously [7-10]. Additional liver stem cells of a 5-month-old subject and an 18-year-old subject were isolated from bulk hepatocytes suspensions (Lonza Walkersville Inc.) as described [7,8] using the same polarization media as described above. The identity of the stem cells isolated were confirmed using the same set of markers as the above using FACS.

Clones of liver stem cells were prepared as follows. Liver stem cells were plated on CellRaft arrays (Cell Microsystems), which contain 12,000 individual portable rafts. Rafts containing single cells were identified under a microscope. The array was then used for culturing the cells. When 8-10 cell clones were generated from single cells, rafts containing these clones were isolated using the CellRaft system and transferred to separate wells of a 96-well plate for culturing and subsequently to 24- 12- / and 6-well plates, followed by 10cm plates, until reaching 1.5 – 3.0 x$10^6$ cells per clone.

Fibroblast clones were prepared as described previously[1].

**Preparing Bulk DNA from bulk cells and clones**

For B lymphocytes, bulk DNA was extracted from PBMCs after depleting all lymphocytes using the DNeasy Blood & Tissue Kit (Qiagen) following the manufacturer's specifications. For hepatocytes, bulk DNA was extracted from total cell suspensions using the same method as for the B lymphocytes. Bulk DNA of fibroblasts was prepared as described previously[1].

**Library preparation and sequencing**

Libraries of the amplified DNA product of single cells, clones and bulk were prepared using the Truseq Nano DNA HT Sample Preparation Kit (Illumina) by Novogene. The libraries were sequenced by Novogene on Illumina HiSeq X Ten platform with 2x150 bp paired-end reads.

**External source of sequencing data**

A part of the single fibroblast sequencing data, including 18 single cells of one human subject and their corresponding bulk DNAs, was obtained from ref[11] (Table S1). The 18 single fibroblasts were amplified using six different whole-genome amplification protocols, i.e., two MDA-based protocols (GE, and Qiagen), Rubicon, GenomePlex, MALBAC and LIANTI.

All single neuron sequencing data, including 36 single neurons from the cerebral cortex of three human subjects and their corresponding bulk DNAs, were obtained from ref[12] (Table S1). The neurons were amplified using MDA.

**Sequence alignment**

Raw sequencing data were quality- and adaptor-trimmed using Cutadapt (v1.8.3)[13] and Trim Galore (v0.4.1) (https://www.bioinformatics.babraham.ac.uk/projects/trim_galore/), and aligned to the human reference genome (hg19) using BWA-MEM (v0.7.12)[14]. PCR duplications were removed using samtools (v0.1.19)[15]. The remaining alignments were subject to indel-realignment using GATK (v3.5)[16] based on indels reported by the 1000 Genomes Project[17], and base-quality recalibration using GATK (v3.5) based on SNPs reported in dbSNP (v138)[18] and the 1000 Genomes project.

**Identifying somatic retrotranspositions**

TraFiC-mem (v2.0) (https://gitlab.com/mobilegenomesgroup/TraFiC) with its default settings was used to identify somatic RTs (L1, Alu, EVRK and others) by comparing alignments of single cells or clones to the alignments of their corresponding bulk[19,20]. In brief, TraFiC used BWA-MEM to search for reads containing retrotransposon-like sequences, and reconstructed insertion break points by *de novo* assembling of the reads identified. The above step was performed for single cells and bulk DNAs separately. Candidate RTs in single cells were then filtered out if they were also found in their corresponding bulk DNAs or found to overlap with known germline retrotransposon polymorphisms reported by the 1000 Genomes Project with the same criteria used previously[19,20]. Neighboring L1 RTs (i.e., distance of insertion sites <1 kb) were combined as a single L1 RT event. Candidate RTs found in cells of different human subjects were filtered out, also to avoid artifacts from germline retrotransposon polymorphisms.

Subtypes of L1 RTs were also identified using TraFiC-mem. They include (a) solo-L1 events, in which either partial or complete LINEs are somatically retrotransposed, (b) partnered transductions, in which a LINE and downstream nonrepetitive sequence are retrotransposed, and (c) orphan transductions, in which only the unique sequence downstream of an active L1 is retro-transposed without the cognate LINE. Based on the transducted nonrepetitive sequences, germline source L1 retrotransposons of RTs were determined by TraFiC-mem.

**PCR validation of variant calling**

PCR primers were designed to validate break points of somatic RTs: the forward primers were complementary to 5’ upstream sequences of the RT break points on the reference genome; the reversed primers covered the RT break points, i.e. part of the primer sequence complementary to the reference genome and the other part complementary to the inserted retrotransposon sequence. PCR was performed for both single cells/clones and their corresponding bulk DNA.

**Permutation test for genomic features**

Gene annotations were obtained from ENSEMBL biomart (GRCh37, release 78). For 5’ and 3’ UTR regions, their 1kb flanking regions were included in the analyses below, because the UTR regions themselves do not cover enough bases to be analyzed. Annotations of CpG islands were obtained from the UCSC genome browser (hg19). CpG island shores were defined as regions within the 2 kb flanking CpG islands.

For each genomic feature, the number of RT insertion sites in the feature was counted using bedtools[21]. Then, regions of the genomic feature were randomly redistributed in the genome using bedtools, and the number of RT insertion sites in the random regions were counted. The above randomization was repeated for 2,000 times. A Monte Carlo P values were determined by comparing the real count with the 2,000 counts obtained from the randomization, as described previously[22].

**Permutation test for TF target regions**

Target regions of 161 transcription factors (TFs) identified using ChIP-sequencing in 91 cell types by the ENCODE project (release 3)[23] were obtained from the UCSC genome browser (hg19). Their 1kb flanking regions were included in the analyses below, because they do not cover enough bases to be analyzed.

For each TF, the number of RT insertion sites in target regions of the TF was counted using bedtools[21]. Then, target regions of the TF were randomly redistributed in the genome using bedtools, and the number of RT insertion sites in the random regions were counted. The above randomization was repeated 2,000 times. A Monte Carlo P value was determined by comparing of the real counts with the 2,000 counts obtained from the randomization, as described previously[22].

**Permutation test for PRC2 target genes**

Four target gene sets of SUZ12, EED, H3K27me3 and PRC2 were obtained from ref[24]. Genes with RT insertions were determined by TraFiC-mem based on ANNOVAR[25]. All the above genes were limited to protein coding genes for the following analyses.

For each target gene set, the number of genes in the intersection of the target gene set (set A) and the set of genes with RT insertions (set B) was counted. Then, two random gene sets, with the same numbers of genes of the set A and B separately, were obtained from the total protein-coding genes, and the number of genes in the intersection of the two random sets was counted. The above randomization was repeated 2,000 times. A Monte Carlo P value was determined by comparing the real count with the 2,000 counts obtained from the randomization, as described previously[22].

**Data availability**

All somatic RTs are provided in Table S4. Raw sequencing data will be available upon publication from dbGAP or SRA database.
