## Supplementary figures and images for "Spontaneous retrotranspositions in normal tissues are rare and associated with cell-type-specific differentiation"

### Supplementary Figures

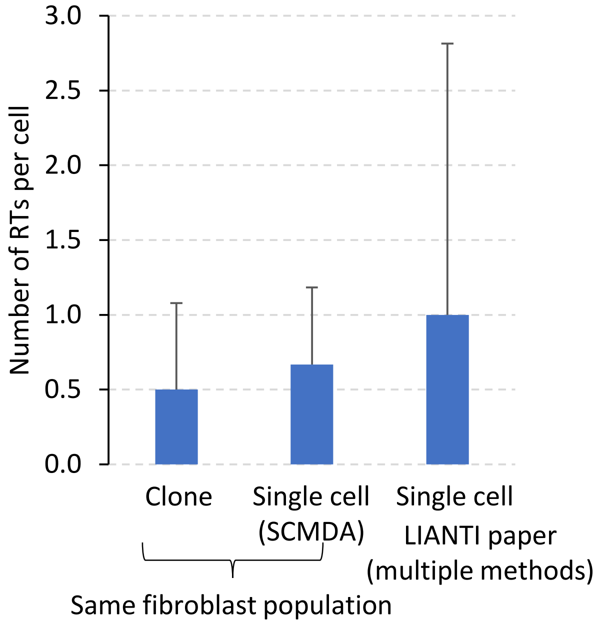

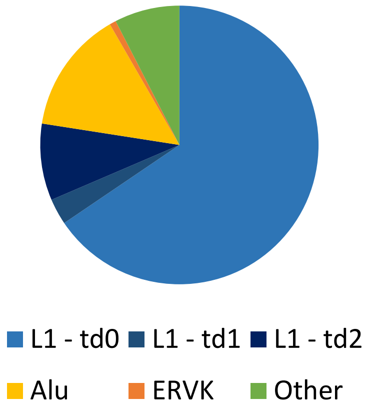

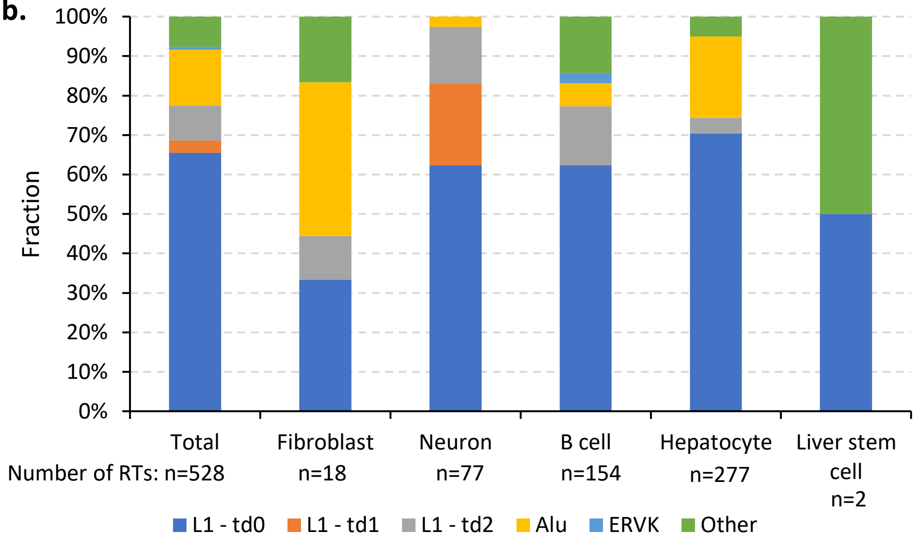

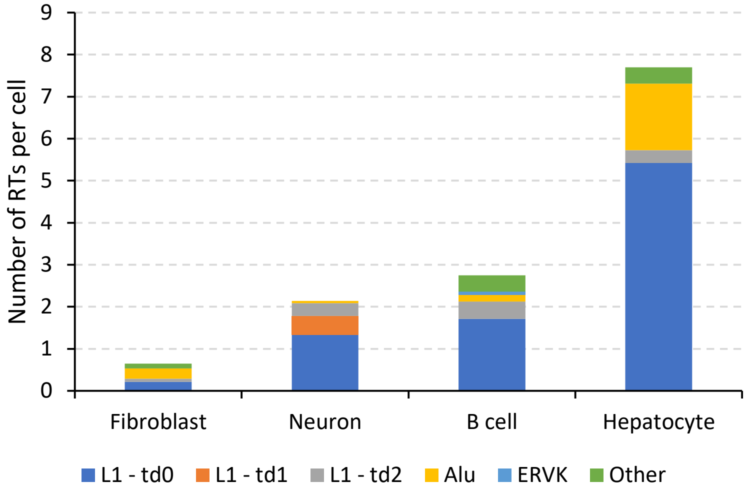

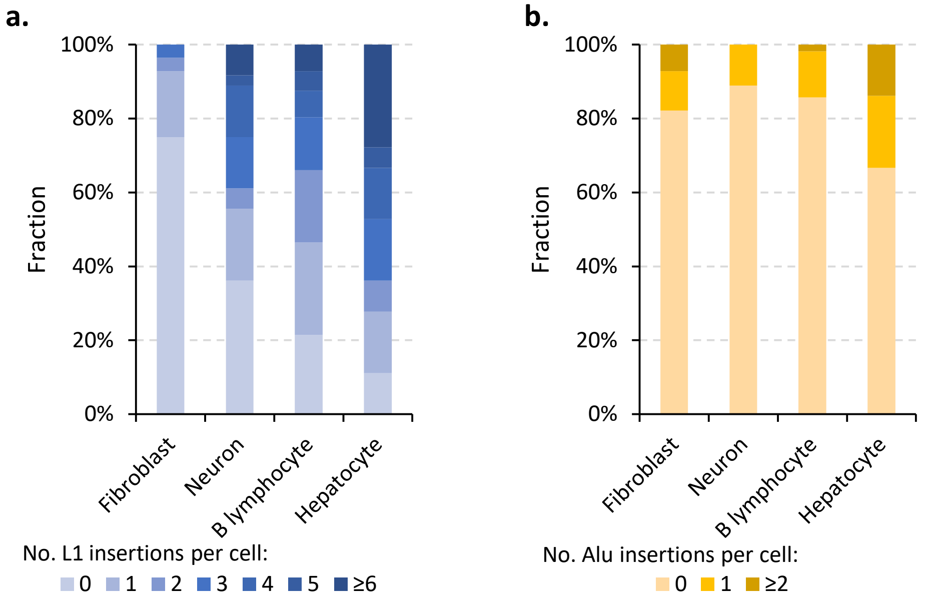

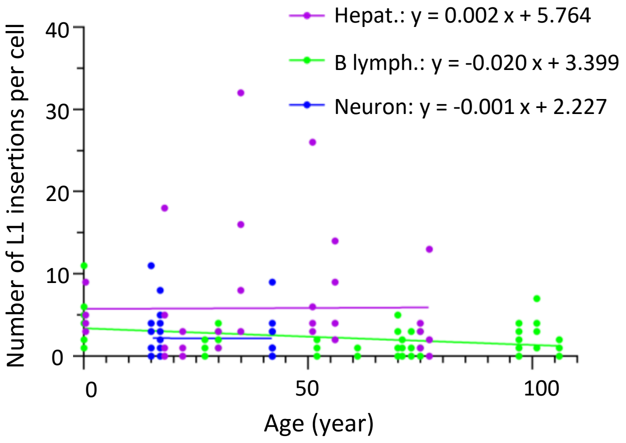

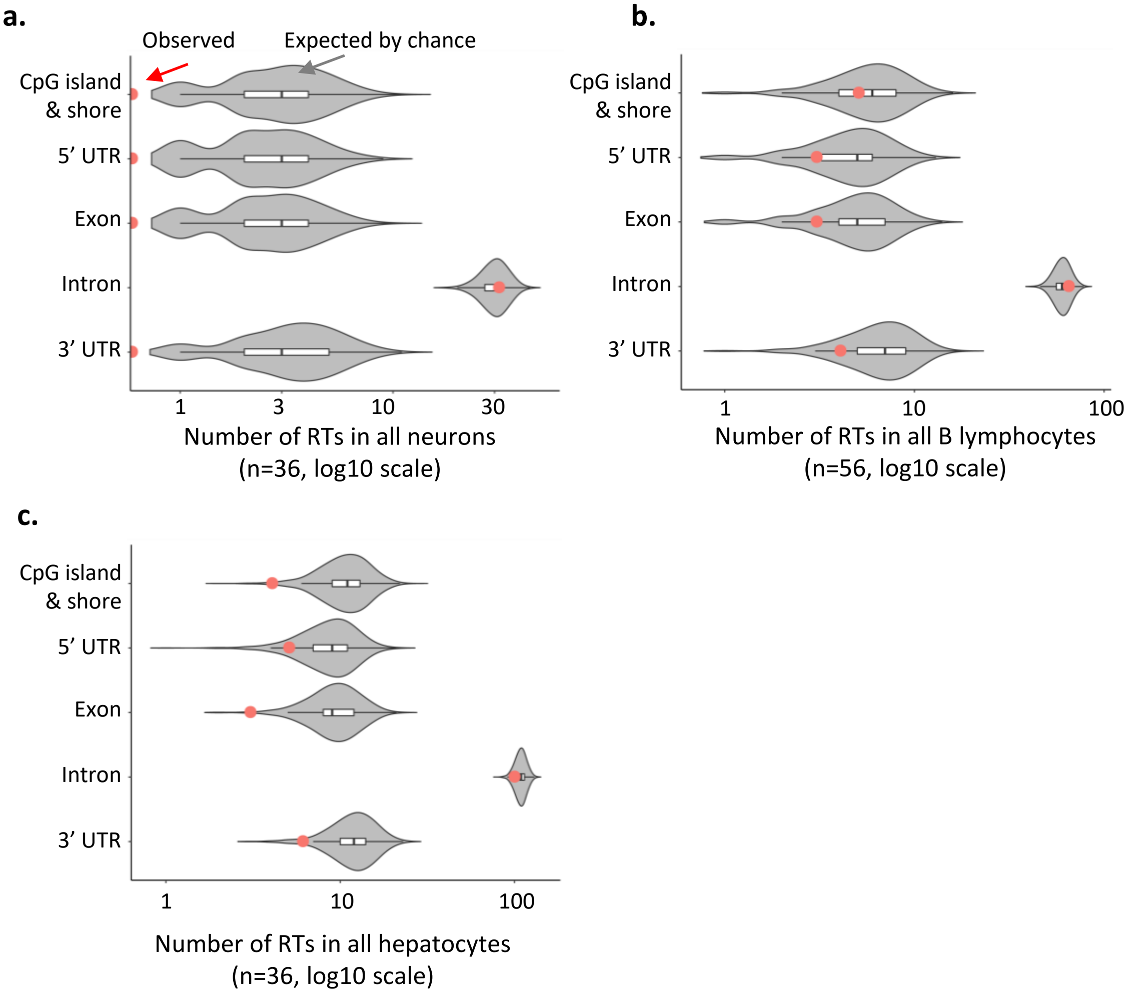

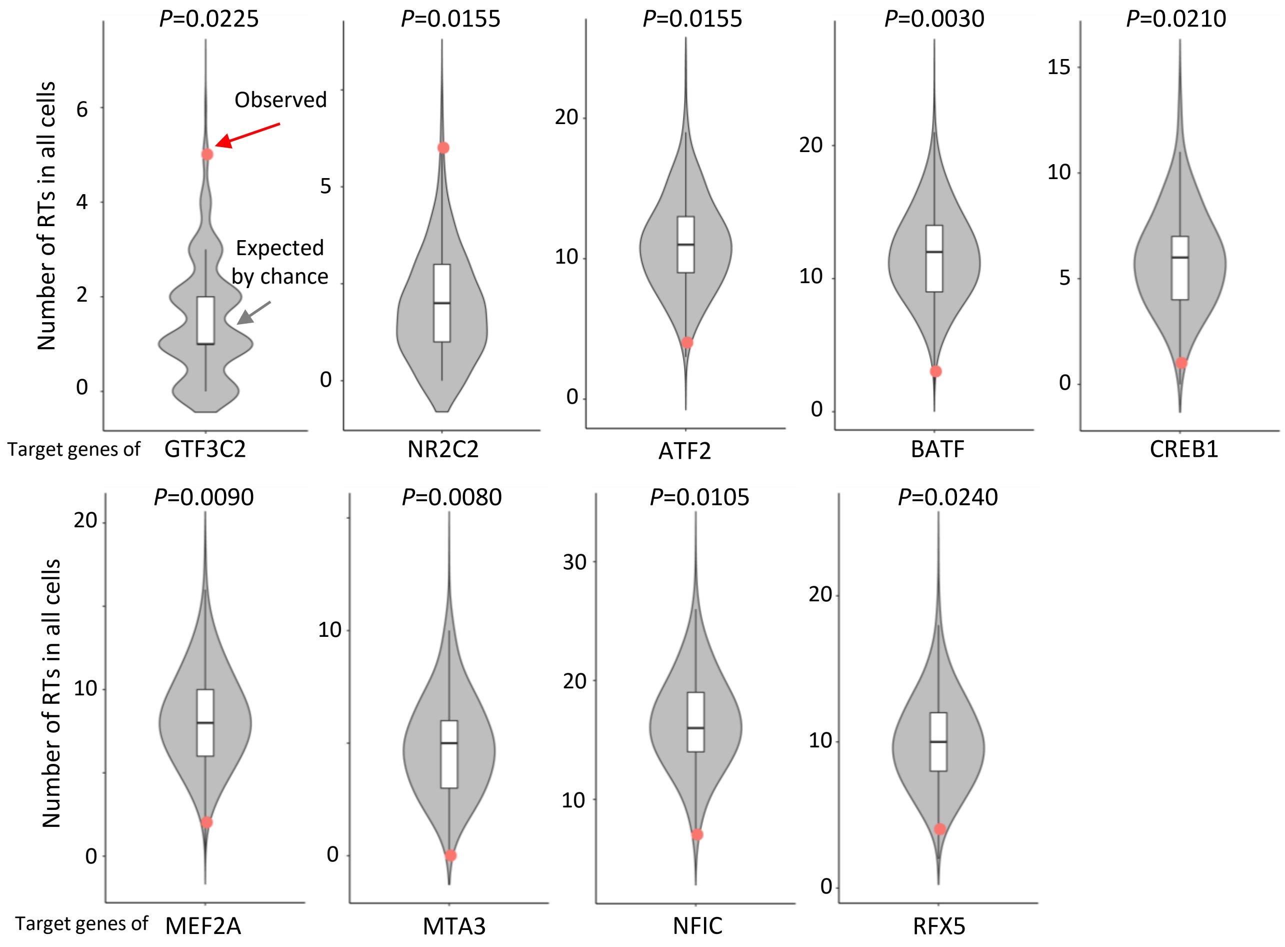

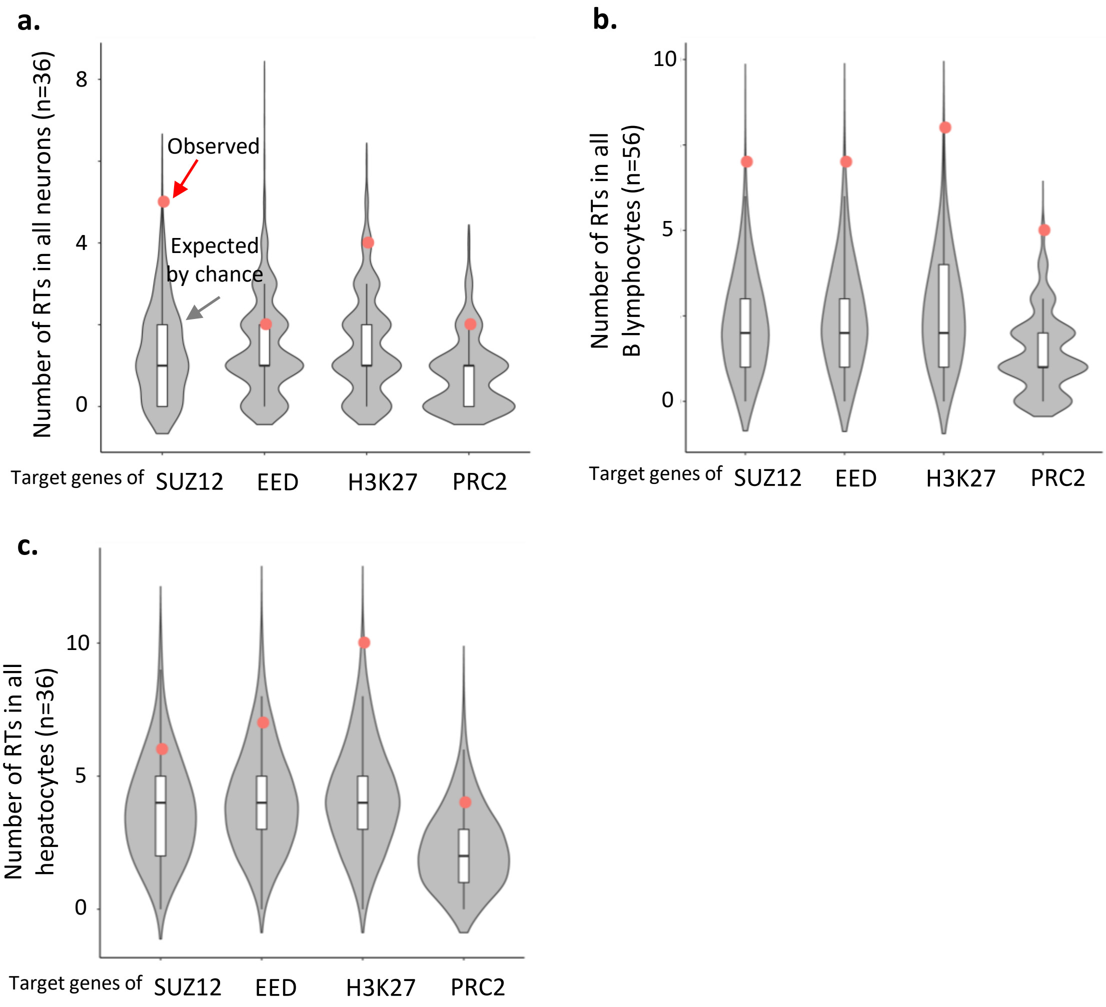

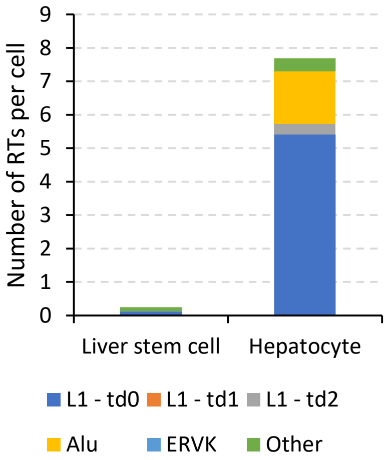
